# 2-hydroxyglutarate-dependent inhibition of TET2 is a molecular link between metabolic reprogramming and smooth muscle cell differentiation

**DOI:** 10.64898/2026.11.18.767744

**Authors:** Jose Emiliano Esparza Pinelo, Alexei Robin, Alexis Richard, Aminat Dosunmu, Jeison Garcia Serrano, Angie Danhi Gonzalez, Hannah Krenz, Jessica Chen, Mario Carrera, Kiara E. Bornes, Shuvra S. Roy, Abhijnan Chatthopadhyay, Dianna M. Milewicz, Eyal Gottlieb, Callie S. Kwartler

## Abstract

**Background:** Smooth muscle cell (SMC) phenotypic modulation is accompanied by metabolic reprogramming, but the mechanisms linking metabolism to contractile gene expression remain unclear. We investigated whether 2-hydroxyglutarate (2-HG), an inhibitor of α-ketoglutarate-dependent dioxygenases, links metabolic changes to regulation of ten-eleven translocation 2 (TET2) in *Acta2*^R179C/+^ SMCs

**Methods:** Primary wild-type (WT) and *Acta2^R^*^179^*^C/+^* mouse aortic SMCs were treated with nicotinamide riboside (NR), exogenous R&L-2-HG, or cholesterol. Endogenous accumulation of L-2HG was achieved by lentivuris-mediated knockdown (KD) of *L2hgdh.* We assessed contractile marker expression, cellular metabolism, TET2 chromatin occupancy, and DNA methylation and hydroxymethylation. L-2HG levels were assessed using liquid chromatography/mass spectroscopy. Carotid artery 5-hydroxymethylcytosine (5hmC) staining was assessed in mice after left carotid artery ligation.

**Results:** *Acta2^R^*^179^*^C/+^* SMCs have elevated 2-HG, reduced TET2 expression and chromatin occupancy, and lower 5hmC enrichment at CArG-containing SMC loci, consistent with repressed TET2 activity. NR boosts oxidative phosphorylation (OXPHOS), reduces 2-HG in WT and mutant SMCs, increases TET2 expression and 5hmC enrichment, and reduces global 5-methylcytosine. Carotid artery 5hmC staining shows changes in the same direction, but group differences are not statistically significant. Exogenous L-2-HG attenuates NR-induced contractile marker expression in *Acta2^R^*^179^*^C/+^* SMCs and also decreases contractile marker expression, reduces TET2 occupancy and 5hmC enrichment, and shifts WT SMC metabolism toward glycolysis, phenocopying the metabolic state of *Acta2*^R179C/+^ SMCs. Endogenous accumulation of L-2HG by *L2hgdh* KD reduces contractile markers in WT SMCS. Cholesterol reduces TET2 and contractile marker expression, decreases 5hmC enrichment, and increases global DNA methylation.

**Conclusions:** These findings support a role for 2-HG in coupling metabolic state to epigenetic changes associated with SMC phenotypic modulation. NR-associated differentiation coincides with increased TET2 expression and DNA hydroxymethylation, identifying metabolic regulation of the SMC epigenome as a potential approach to preserving contractile gene expression.

## Introduction

Smooth muscle cell (SMC) differentiation is a metabolically regulated process essential for maintaining vascular tone and function^1–3^. Unlike terminally differentiated skeletal or cardiac muscle cells, SMCs retain phenotypic plasticity, and the phenotypic modulation of SMCs is a contributor to cardiovascular diseases such as atherosclerosis and thoracic aortic dissections^4–6^. Modulated SMCs lose their contractile identity and often exhibit features of cancer cells^7^ including increased proliferation, migration, and a shift from oxidative phosphorylation (OXPHOS) toward glycolysis as a source of cellular energy^4,8^. Metabolic intermediates can directly influence gene expression by acting as cofactors or competitive inhibitors of chromatin-modifying enzymes, providing a mechanistic link between metabolic state and cell identity^1,3,4,910^. However, whether specific metabolic intermediates drive SMC phenotypic switching through defined epigenetic mechanisms remains largely unexplored.

*ACTA2* p.Arg179 (R179) pathogenic variants cause multisystemic smooth muscle dysfunction syndrome (MSMDS), a severe congenital vasculopathy in which SMC dysfunction manifests across multiple organ systems, including moyamoya disease (MMD)-like cerebrovascular occlusions, early-onset aortic aneurysms and dissections, and widespread visceral hypomotility^5,11–15^. Our previous work showed that *ACTA2* p.R179C SMCs are both incompletely differentiated and metabolically rewired, providing a model to examine whether specific metabolic intermediates directly drive epigenetic changes associated with SMC differentiation^16,17^. Specifically, explanted mouse *Acta2*^R179C/+^ SMCs exhibit a phenotype with compromised contractile gene expression, increased migration, and a pathological reliance on glycolysis over OXPHOS for energy production^17^. As a result of these SMC phenotypic changes, conditional knock-in mice (*Acta2*^SMC-R179C/+^) develop MMD-like carotid lesions and stroke-associated mortality following left carotid artery ligation (LCAL) injury^17^. Remarkably, both cellular and *in vivo* phenotypes are rescued by nicotinamide riboside (NR), an NAD^+^ precursor that boosts electron transport chain activity and OXPHOS^18^. In doing so, NR reduces the accumulation of glycolytic and tricarboxylic acid (TCA) cycle intermediates, including 2-hydroxyglutarate (2-HG), while simultaneously restoring contractile marker expression and attenuating MMD-like disease in the *Acta2*^SMC-R179C/+^ mice^17^. However, the mechanism by which this metabolic reprogramming leads to transcriptional rescue of SMC differentiation was not clear from prior work.

Among the intermediates reduced by NR, 2-HG stands out as a signaling metabolite capable of reshaping the epigenetic landscape, making it a strong candidate to link the metabolic alteration of *Acta2*^R179C/+^ SMCs with contractile differentiation. Mechanistically, 2-HG is a competitive inhibitor of α-ketoglutarate (α-KG)-dependent dioxygenases, including ten-eleven translocation 2 (TET2)^19^, a DNA dioxygenase that catalyzes the oxidation of 5-methylcytosine (5mC) to 5-hydroxymethylcytosine (5hmC) and has been established as a master epigenetic regulator of SMC contractile gene expression^20–23^. We tested the hypothesis that 2-HG accumulation in glycolytic *Acta2*^R179C/+^ SMCs inhibits TET2, and that NR rescues SMC differentiation by reducing 2-HG and restoring TET2-dependent demethylation of contractile gene promoters. Here we show that *Acta2*^R179C/+^ SMCs have elevated 2-HG and reduced TET2 levels, chromatin occupancy, and 5hmC enrichment at CArG-containing SMC loci. NR treatment increases TET2 expression and DNA hydroxymethylation in cultured SMCs, coinciding with increased contractile marker expression. To determine whether this mechanism affects SMC phenotype beyond the *ACTA2* p.R179C context, we show that exogenous addition of 2-HG is sufficient to dedifferentiate WT SMCs, inducing a metabolic shift toward glycolysis, reduced expression of contractile markers, increases in DNA methylation, and decreases in DNA hydroxymethylation that recapitulate features of *Acta2*^R179C/+^ SMCs. Finally, treatment with methyl-β-cyclodextrin complexed cholesterol (MBD-Chol) is known to induce SMC phenotypic modulation in culture and also to drive glycolytic metabolism^24^. Here, we show that MBD-Chol reduces TET2 expression and 5hmC enrichment, extending the association between altered DNA hydroxymethylation and SMC phenotypic modulation to a second model. Together, these findings support a link between 2-HG, TET2 regulation, and SMC differentiation, and identify a potential route by which metabolic changes affect contractile gene expression.

## Materials and Methods

### Data Availability

The authors declare that all data supporting the findings of this study are available within the article and its Supplemental Material, or from the corresponding author upon reasonable request. The stable isotope tracing and 2-hydroxyglutarate quantification data referenced in this study were previously published by Kaw et al. ^17^ and are available in that article and its associated Source Data. This study did not generate large-scale sequencing or other datasets requiring deposition in a public repository. Antibodies, primer sequences, cell models, and other key reagents are listed in the Major Resources Table.

### Cell culture and metabolic treatments

Primary smooth muscle cells (SMCs) were explanted from the ascending aortas of 8-week-old, age- and sex-matched wild-type and *Acta2*^SMC-R179C/+^ mice and cultured as previously described^17,25^. Briefly, ascending aortic tissue pooled from 3-5 mice per genotype was cleaned, digested overnight in collagenase type I, elastase, and soybean trypsin inhibitor, and cultured in Smooth Muscle Basal Medium (SmBm; PromoCell) supplemented with 20% FBS, insulin, epidermal growth factor, fibroblast growth factor, HEPES, sodium pyruvate, L-glutamine, and antibiotic/antimycotic. Explanted *Acta2*^SMC-R179C/+^ SMCs (designated *Acta2*^R179C/+^ SMCs) are uniformly heterozygous for the *Acta2* R179C variant¹⁷. All experiments were performed with cells at passages 2-5, and WT and *Acta2*^R179C/+^ SMCs were passage-matched for each independent experimental replicate.

For nicotinamide riboside (NR) treatment, SMCs were treated with 0.25 mg/mL NR triflate (Carbosynth Biosynth) or an equivalent volume of ddH₂O (vehicle) for 5 days, with NR- or vehicle-containing medium replenished on days 1, 3, and 5, as previously described¹⁷. Cells were then serum-starved for 24 h in SmBm containing 1% FBS with NR or vehicle prior to harvest, unless otherwise noted.

For L-2-HG and D-2-HG treatments, SMCs were treated with 10, 50, or 100 nM L- or D-α-hydroxyglutaric acid disodium salt (L-2-HG or D-2-HG; Cayman Chemical, Item No. 11876) or an equivalent volume of PBS (vehicle) for 72 h in treatment medium, consisting of SmBm supplemented with 1% FBS, HEPES, sodium pyruvate, L-glutamine, and antibiotic/antimycotic.

For NR and L-2-HG co-treatment, SMCs were treated concurrently with 0.25 mg/mL NR and 50 nM L-2-HG following the same 5-day schedule used for NR treatment, with medium containing both compounds replenished on days 1, 3, and 5, followed by 24 h of serum starvation in SmBm containing 1% FBS with both compounds prior to harvest.

WT SMCs were treated with 10 µg/mL water-soluble cholesterol (free cholesterol complexed to methyl-β-cyclodextrin, MBD-Chol; Millipore Sigma) or vehicle (deionized water) for 72 h, as previously described^24^. MBD-Chol contains approximately 40 mg cholesterol per gram of product, and all treatment concentrations were based on cholesterol weight. Treatments were performed in high-glucose Dulbecco’s Modified Eagle Medium (DMEM; Cellgro) supplemented with 10% FBS (Gibco), 1% antibiotic/antimycotic (Millipore Sigma), and 0.2% bovine serum albumin (Fisher Scientific) at 37 °C and 5% CO₂.

### Immunoblotting

SMCs were treated as indicated and lysed in RIPA buffer (50 mM Tris-HCl pH 7.4, 1% NP-40, 0.25% Na-deoxycholate, 150 mM NaCl, 1 mM EDTA, 1 mM PMSF) supplemented with protease and phosphatase inhibitor cocktails (Millipore Sigma), as previously described¹⁷. Lysates were sonicated, and protein concentration was quantified by Bradford assay (Bio-Rad). Equivalent amounts of protein (3–30 µg, depending on the target) were boiled in loading buffer, separated on 4–20% TGX SDS-PAGE gels (Bio-Rad), and transferred to PVDF membranes (Bio-Rad Trans-Blot Turbo). Membranes were blocked with 5% non-fat dry milk or 5% BSA in TBST for 1 h and incubated overnight at 4 °C with primary antibodies (1:1000), followed by the corresponding HRP-conjugated secondary antibodies (1:3000) for 1 h at room temperature. Histone H3 was used as a loading control, as GAPDH is a glycolytic enzyme whose expression may be altered in mutant and metabolically manipulated SMCs. Bands were visualized by chemiluminescent substrate (Bio-Rad) on a Bio-Rad ChemiDoc imaging system, and band intensities were quantified using Image Lab software (Bio-Rad). Antibodies are listed in the Major Resources Table.

### Quantitative real-time PCR (RT-qPCR)

Total RNA was isolated from treated SMCs using the PureLink RNA kit (Thermo Fisher Scientific) and quantified by NanoDrop (Thermo Fisher Scientific), as previously described¹⁷. 50 ng of RNA per sample was reverse transcribed using qScript cDNA synthesis reagent (Vazyme). Quantitative real-time PCR was performed using SYBR Green chemistry (Millipore Sigma) or TaqMan probes (Applied Biosystems) with the appropriate master mixes (Vazyme), with TaqMan chemistry used for contractile gene transcripts and SYBR Green for all other genes. Reactions were performed in triplicate on a CFX Real-Time PCR instrument (Bio-Rad), and relative gene expression was calculated using the ΔΔCT method with *18S* as the endogenous control. Primer sequences and TaqMan probe identifiers are listed in the Major Resources Table.

### *In vivo* immunofluorescence

All animal studies were performed according to protocols approved by the Institutional Animal Care and Use Committee at the University of Texas Health Science Center at Houston and in accordance with the National Institutes of Health guidelines on the care and safety of laboratory animals (Animal protocol number AWC-24-0122). At 8 weeks of age, female WT and Acta2SMC-R179C/+ mice were randomized into treatment groups of either saline or NR chloride. For NR treatment, NR was dissolved in saline and administered through intraperitoneal injection at a final dose of 1000 mg/kg body weight every other day over a 26-day treatment period beginning 5 days before left carotid artery ligation (LCAL)^17^. During the surgery, mice were anesthetized with isoflurane, bupivacaine analgesia was administered via subcutaneous injection, and the neck region was shaved with Nair and sanitized with alcohol. A 3 mm midline incision was made in the neck with sterilized scissors to expose the left common carotid artery, which was dissected from the carotid sheath and vagus nerve, then ligated with a 5-0 silk suture at the bifurcation site where the ICAs separate from the external carotid artery. Open skin was closed by stitches with 5-0 suture, and mice were allowed to recover^17^. At 21 days after ligation, animals received an intraperitoneal injection of Avertin (2.5%, 350 mg/kg, terminal dose), followed by exsanguination and perfusion fixation with 20 mL of 1× PBS (pH 7.4), followed by 20 mL of 10% neutral-buffered formalin for 5 min through the left ventricle under physiological pressure. The left and right carotid arteries were harvested, further fixed in 10% neutral-buffered formalin overnight at room temperature, dehydrated, and embedded in paraffin. Blocks were sectioned at 5 µm.

Formalin-fixed, paraffin-embedded sections were stained by immunofluorescence with primary antibodies for SMC-specific α-actin (α-SMA, Abcam #ab21027) and 5hmC (ActiveMotif #39999). Briefly, after deparaffinization and rehydration, carotid sections were processed for antigen retrieval using sodium citrate buffer (10 mM sodium citrate, 0.05% Tween 20, pH 6.0) at 95 °C for 30 minutes, cooled for 30 min, permeabilized with PBS-0.3% Triton X-100, then blocked in 5% bovine serum albumin in PBS-0.1% Triton X-100 for 1 hour at room temperature, followed by primary antibody (diluted at 1:100 in PBS-0.1% Triton X-100) incubation at 4 °C overnight. Tissue sections were incubated with secondary antibody (Thermo Fisher # A-11055, A-11005) at room temperature for 1 h (diluted at 1:500 in PBS-0.1% Triton X-100), then mounted with 4′,6-diamidino-2-phenylindole (DAPI) (VECTASHIELD Anti-fade Mounting Medium, H-1200, VectorLabs). Immunofluorescent images were obtained using a Leica DMi8 confocal microscope.

FIJI software was used to quantify nuclear 5hmC in medial SMCs (one section per mouse, the whole aortic ring was quantified)^26^. A mask was created using α-SMA to delineate the aortic medial layer, then medial DAPI stains were used to confirm nuclear area via selection and merging into a singular region of interest for quantification. That merged region of interest was applied to the 5hmC channel and each individual nucleus was counted as positive or negative for 5hmC signal. The percentage of 5hmC+ nuclei on each section was graphed in Figure 3B.

### Chromatin Immunoprecipitation

Cells were crosslinked with formaldehyde for 10 minutes at room temperature, followed by quenching with glycine for 5 minutes. Cells were then washed twice with ice-cold PBS, pelleted, and stored at −80 °C until further processing. ChIP lysates were prepared using the SimpleChIP Magnetic Separation Kit (Cell Signaling Technology) following the manufacturer’s protocol and then incubated overnight at 4 °C with rotation in 1× ChIP buffer with either the target TET2 antibody (Abcam ab309481) or a species-matched IgG control (Cell Signaling Technology #2729S). Magnetic protein A/G beads were added to the samples and incubated for 4 hours at 4 °C to capture antibody-bound chromatin complexes. The beads were then washed three times with low-salt wash buffer and once with high-salt wash buffer at 4 °C to reduce nonspecific binding. Chromatin was eluted from the beads in ChIP elution buffer at 65 °C for 30 minutes while shaking. Crosslinks were reversed by incubation at 65 °C with salt and proteinase K. DNA was purified and analyzed by quantitative PCR (qPCR). For each sample, 2% of the total chromatin input was reserved prior to immunoprecipitation for normalization. ChIP-qPCR enrichment was calculated relative to input, with species-matched IgG immunoprecipitation used as a negative control.

### Hydorxymethylated DNA Immunoprecipitation

Genomic DNA was extracted from treated WT and *Acta2*^R179C/+^ SMCs using the DNeasy Blood & Tissue kit (Qiagen) and sheared to fragments of approximately 300–500 bp using a Branson probe sonicator (6 cycles of 15 s at 60% amplitude), with fragment size verified prior to immunoprecipitation. hMeDIP was performed using the Diagenode hMeDIP kit according to the manufacturer’s protocol. Briefly, fragmented DNA was denatured and immunoprecipitated with the kit’s monoclonal anti-5-hydroxymethylcytosine (5hmC) antibody or a species-matched IgG negative control, and antibody-bound DNA was captured with DiaMag protein G–coated magnetic beads. A fraction of fragmented DNA was reserved as input prior to immunoprecipitation, and 5hmC and unmethylated spike-in controls included in the kit were used to confirm immunoprecipitation specificity. Recovered DNA was purified and analyzed by qPCR at CArG-containing promoter regions of SMC contractile genes (*Myh11*, *Acta2*, *Cnn1*, *Tagln*, *Dhx32os*, *Tspan2*). Enrichment was calculated as percent recovery relative to input [(IP / input) × 100]. Primer sequences are listed in the Major Resources Table.

### Global DNA Methylation Quantification

Global 5-methylcytosine (5mC) DNA methylation levels were quantified using the MethylFlash Methylated DNA Quantification Kit (EpigenTek, Farmingdale, NY) in a 48-well ELISA-based colorimetric format. Genomic DNA was extracted from WT and *Acta2^R^*^179^*^C/+^* SMCs treated with NR or L-2-HG as indicated using the DNeasy Blood & Tissue Kit (Qiagen), and DNA concentration was determined by NanoDrop spectrophotometry. Samples were diluted to 200 ng per reaction, consistent with the manufacturer’s recommended optimal input for efficient DNA binding to the strip wells.

A standard curve was prepared using the provided methylated DNA standards at concentrations of 0.5, 1.0, 1.2, 1.5, 2.0, 5.0, and 10.0 ng/µL. DNA samples were added directly to the high-affinity DNA-binding microplate wells. Following binding, methylated DNA was detected using a capture antibody specific for 5mC, followed by a detection antibody and signal enhancer solution, according to the manufacturer’s protocol. Absorbance was measured at 450 nm using a microplate reader, and the absolute amount of methylated DNA in each sample was calculated from the standard curve using the formula provided in the manufacturer’s instructions.

### Extracellular flux Analysis

Extracellular flux analysis was performed using an Agilent Seahorse XF Flex analyzer (Seahorse Biosciences) to assess mitochondrial respiration via oxygen consumption rate (OCR) and glycolysis via extracellular acidification rate (ECAR). WT and *Acta2*^R179C/+^ SMCs were treated as indicated and seeded at 100,000 cells/well in SmBm supplemented with 20% FBS, insulin, epidermal growth factor, fibroblast growth factor, HEPES, sodium pyruvate, L-glutamine, and antibiotic/antimycotic overnight. OCR was measured using the Mito Stress Test Kit and ECAR was measured using the Glycolysis Stress Test Kit (Agilent), following the manufacturer’s protocols. Assays were performed with 5–10 technical replicates per condition, and graphs are representative of at least three independent experiments.

### Lentivirus infection for L2HGDH shRNA Knockdown

Lentiviral vectors for L2HGDH knockdown were generated using MISSION shRNA constructs targeting mouse *L2hgdh* (Sigma-Aldrich; clone IDs: TRCN0000173393, TRCN0000173556, TRCN0000175075, TRCN0000322334, and TRCN0000322335) A non-targeting shRNA (shCtrl) served as a negative control. Lentivirus was produced in HEK293T cells using shRNA constructs along with packaging plasmids pMD2G and psPax2 (Addgene). Viral medium was filtered and concentrated with Lenti-X Concentrator and added to WT SMCs SmBm media. Knockdown efficiency was assessed by RT-qPCR, and clones achieving greater than 50% reduction in *L2hgdh* expression were selected for downstream experiments. Endogenous L-2-HG accumulation following *L2hgdh* knockdown was confirmed by chiral LC-MS/MS.

### Statistical Analysis

All statistical analyses were performed using GraphPad Prism (version 11.1.0; GraphPad Software). Data are presented as mean ± standard deviation (SD). Comparisons between two groups were analyzed by two-tailed unpaired Student’s t-test. Comparisons of a single variable across more than two groups were analyzed by one-way ANOVA with the appropriate multiple comparisons test; for dose-response experiments and shRNA knockdown validation, each condition was compared to the untreated control using Dunnett’s multiple comparisons test. Experiments assessing two variables (genotype and treatment) were analyzed by two-way ANOVA with Tukey’s multiple comparisons test. In vivo 5hmC immunostaining was quantified by one-way ANOVA with Tukey’s multiple comparisons test across the three groups (WT, *Acta2*^SMC-R179C/+^, and NR-treated *Acta2*^SMC-R179C/+^). A p-value of P < 0.05 was considered statistically significant. The specific statistical test, number of independent experiments, and definition of replicates for each experiment are indicated in the corresponding figure legends.

## Results

### Increasing OXPHOS in *Acta2*^R179C/+^ SMCs reduces 2-HG and rescues *TET2* expression and chromatin occupancy

*Acta2*^R179C/+^ SMCs show significantly reduced expression and protein accumulation of TET2 (Figure 1A, S1A), consistent with a disrupted regulatory mechanism leading to incomplete SMC differentiation and disease progression^22,27^. Liquid chromatography-tandem mass spectrometry using isotope labeling revealed higher relative 2-HG levels in *Acta2*^R179C/+^ SMCs compared to WT SMCs, and treatment with NR significantly reduced 2-HG accumulation in *Acta2*^R179C/+^ SMCs but failed to do so in WT SMCS (Figure 1B)^17,28^. Consistent with an increase in SMC differentiation, treatment of both WT and *Acta2*^R179C/+^ SMCs with NR upregulates TET2 at both the mRNA and protein level (Figure 1C, S1B). Chromatin immunoprecipitation (ChIP) showed reduced TET2 occupancy at CArG-containing promoter regions of *Myh11*, *Acta2*, *Cnn1*, *Tagln*, and *Tspan2* in *Acta2^R^*^179^*^C/+^* SMCs; the genotype difference at *Dhx32os* was not statistically significant. NR increased TET2 occupancy at *Acta2*, *Myh11*, and *Dhx32os*. Changes at *Cnn1* and *Tspan2* were not statistically significant, and increased occupancy was not observed at *Tagln* (Figure 1D).

**Fig. 1:**
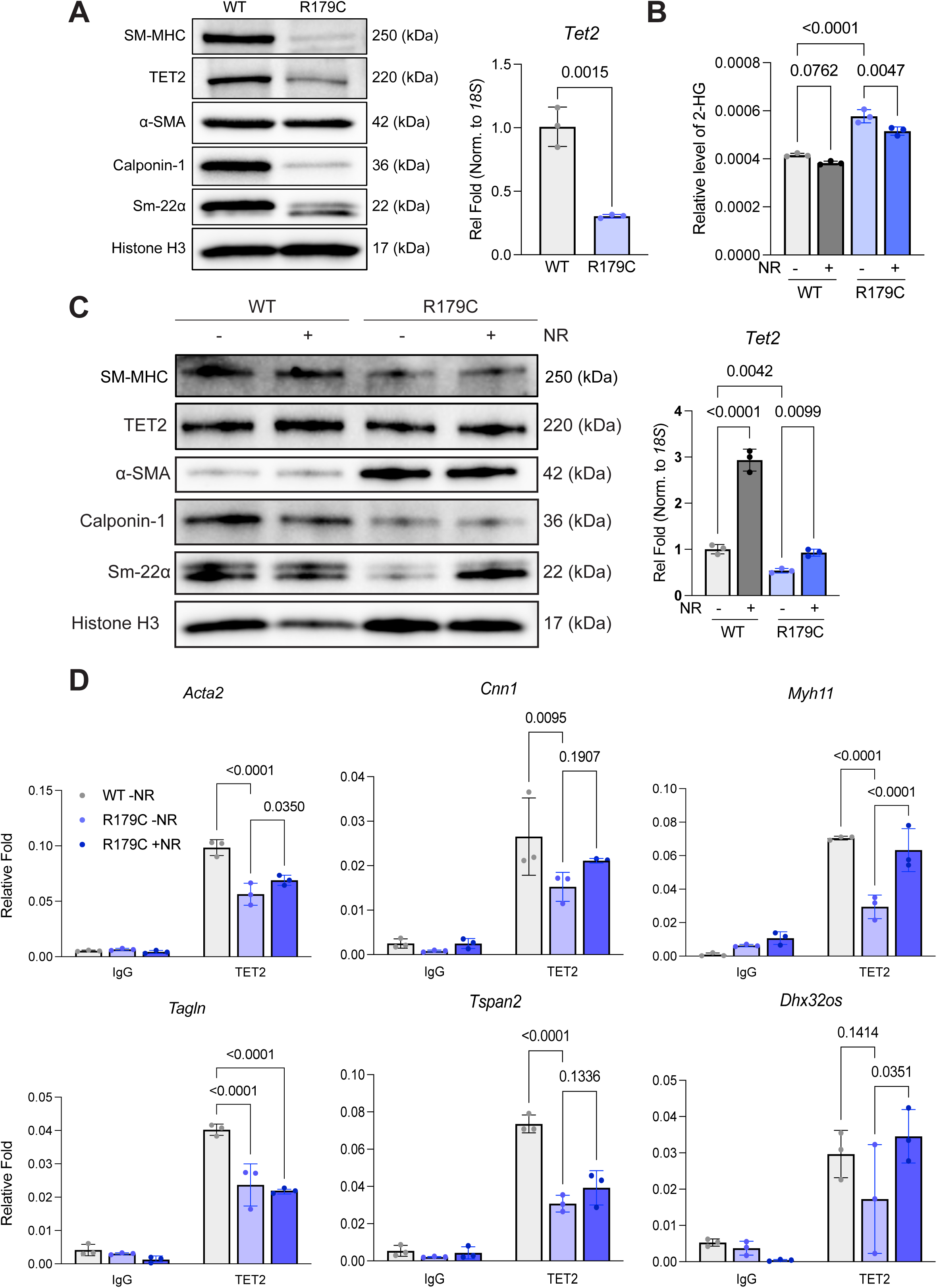
*Acta2*^R179C/+^ mouse smooth muscle cells have decreased TET2 levels rescued by a shift in OXPHOS. **A,** Immunoblot analysis demonstrates reduced levels of TET2 in *Acta2*^R179C/+^ SMCs (left), with a significant reduction in gene expression assessed through RT-qPCR (right). Each data point represents a technical replicate (n=3), and results are representative of more than eight independent experiments. **B,** Relative levels of 2-HG in WT and in *Acta2*^R179C/+^ SMCs treated with or without NR, as quantified by liquid chromatography-tandem mass spectrometry (LC-MS/MS) using 1,2-¹³C₂-glucose isotope tracing. *Acta2*^R179C/+^ SMCs have elevated 2-HG compared to WT, and NR reduces 2-HG in mutant SMCs; the change in WT SMCs is not statistically significant. Each data point represents an independent experiment (n=3). **C,** Immunoblot analysis demonstrates that nicotinamide riboside (NR) increases levels of TET2 in *Acta2*^R179C/+^ SMCs (left), with a significant increase in expression assessed through RT-qPCR (right). Each data point represents a technical replicate (n=3), and results are representative of more than eight independent experiments. **D,** Chromatin immunoprecipitation (ChIP) analysis confirms decreased occupancy of TET2 across multiple SMC contractile gene promoters in *Acta2*^R179C/+^ SMCs, with occupancy increased by NR treatment. Species-matched IgG pulldown is shown as a negative control for each pulldown. Each data point represents a technical replicate (n=3), and results are representative of more than three independent experiments. Statistical significance was assessed for two groups (A) using an unpaired two-tailed t-test with Welch’s correction and for three or more groups (B-D) by two-way ANOVA with Tukey’s post hoc correction for multiple comparisons. Quantifications of independent immunoblot replicates can be found in Supplemental Figure 1A,B. P-values are indicated above brackets. All data are presented as mean ± standard deviation. SMC, smooth muscle cell; TET2, ten-eleven translocation 2; WT, wild-type; R179C, *Acta2*^R179C/+^; NR, nicotinamide riboside; SM-MHC, smooth muscle myosin heavy chain; α-SMA, α-smooth muscle actin.

### Global and site-specific activity of TET2 is rescued by increasing OXPHOS in *Acta2*^R179C/+^ SMCs

Hydroxymethylated DNA Immunoprecipitation (hMeDIP) shows significantly decreased global 5hmC enrichment in *Acta2*^R179C/+^ SMCs compared to WT SMCs. Treatment with NR significantly increases 5hmC enrichment globally in both genotypes, especially in *Acta2*^R179C/+^ SMCs, indicating a rescue of TET2 activity (Figure 2A). Conversely, global DNA methylation was assessed by measuring levels of 5mC using MethylFlash™ Global DNA Methylation (5-mC) ELISA Easy Kit*. Acta2*^R179C/+^ SMCs present with significantly higher global 5mC enrichment compared to WT SMCs. Treatment with NR significantly decreases the amount of 5mC globally in *Acta2*^R179C/+^ SMCs, correlating to the increase in global 5hmC levels (Figure 2B, S2A). Loss of locus-specific enrichment of 5hmC at CArG-containing promoter regions of key SMC genes (*Myh11*, *Acta2*, *Cnn1*, *Tagln*, *Dhx32os*, and *Tspan2)* was observed in *Acta2*^R179C/+^ SMCs compared to WT. CArG 5hmC enrichment was significantly increased at all tested promoter sites following NR treatment in *Acta2*^R179C/+^ SMCs (Figure 2C). Together, these results show that NR-mediated metabolic reprogramming toward OXPHOS restores TET2 activity at contractile gene promoters, driving the conversion of 5mC to 5hmC and reactivating the SMC contractile program.

**Fig. 2:**
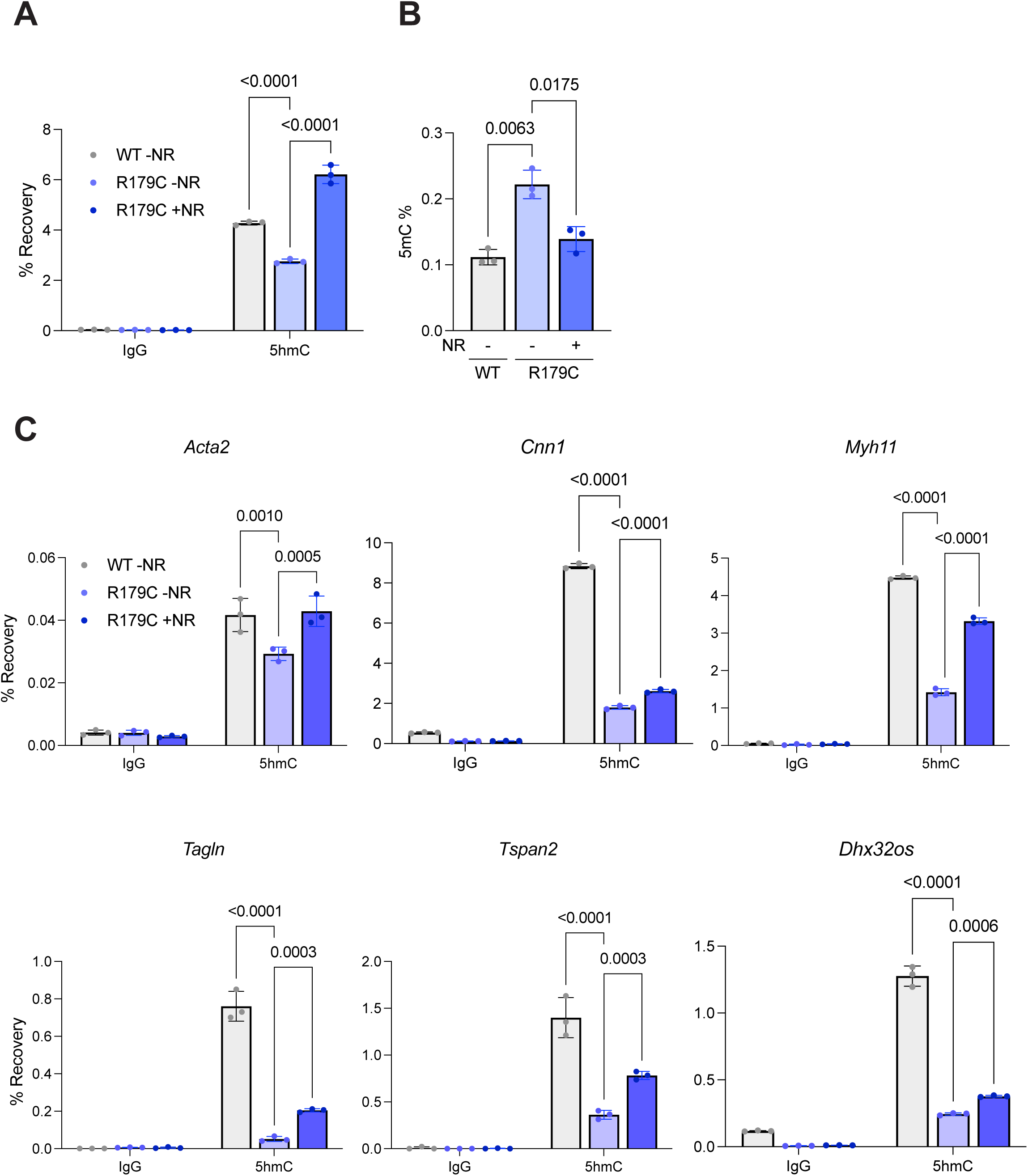
*Acta2*^R179C/+^ mouse smooth muscle cells have decreased TET2 activity rescued by a shift in OXPHOS. **A,** Hydroxymethylated DNA immunoprecipitation (hMeDIP) analysis shows reduced recovery of 5-hydroxymethylcytosine (5hmC) in *Acta2*^R179C/+^ SMCs, with levels increased by NR treatment. Species-matched IgG pulldown is shown as a negative control for each pulldown. **B,** Levels of 5-methylcytosine (5mC) are increased in *Acta2*^R179C/+^ SMCs by MethylFlash ELISA analysis, and 5mC is reduced by NR treatment. **C,** hMeDIP confirms decreased levels of 5hmC across multiple SMC contractile gene promoters in *Acta2*^R179C/+^ SMCs, with levels increased by NR treatment. Species-matched IgG pulldown is shown as a negative control for each pulldown. For all graphs in this figure, each data point represents a technical replicate (n=3), and results are representative of more than three independent experiments. Statistical significance was assessed by two-way ANOVA with Tukey’s post hoc correction for multiple comparisons. P-values are indicated above brackets. All data are presented as mean ± standard deviation. SMC, smooth muscle cell; TET2, ten-eleven translocation 2; WT, wild-type; R179C, *Acta2*^R179C/+^; NR, nicotinamide riboside; 5hmC, 5-hydroxymethylcytosine; 5mC, 5-methylcytosine.

### NR increases TET2 activity in LCAL-induced MMD-like occlusive lesions in *Acta2^SMC-^*^R179C/+^ mice

Generally, high TET2 activity in vascular lesions sites indicates a protective, anti-inflammatory and stabilizing response to injury^27,29^. TET2 stabilization prevents excessive inflammation and cellular dysfunction in SMCs, enabling proper functioning of SMCs and efficient resolution of injury^30^. To examine whether the changes in DNA hydroxymethylation observed in cultured SMCs extend to arterial tissue, we assessed 5hmC levels in medial SMCs from WT and *Acta2*^SMC-R179C/+^ mice subjected to left carotid artery ligation (LCAL). To compare medial SMCs between genotypes, we examined the contralateral uninjured right carotid artery at postoperative day (POD) 21, which shows no lesion in either genotype. This avoids the neointimal lesions and cellular heterogeneity present in the injured left carotid, where *Acta2*^SMC-R179C/+^ mice develop unresolved occlusive neointima-thrombus that is rescued by NR treatment. Representative images suggested lower medial 5hmC (delineated by dotted white line) staining in mutant arteries and higher staining after NR treatment (Figure 3A). Quantification showed a lower proportion of 5hmC-positive medial SMCs in untreated *Acta2^SMC-R^*^179^*^C/+^* mice than in WT mice (P=0.0801), and a higher proportion in NR-treated than in untreated mutant mice (P=0.0850; Figure 3B). Neither comparison reached statistical significance. These observations are consistent in direction with the cultured-cell findings but do not establish an *in vivo* effect of NR on TET2 activity in *Acta2*^SMC-R179C/+^ mice.

**Fig. 3:**
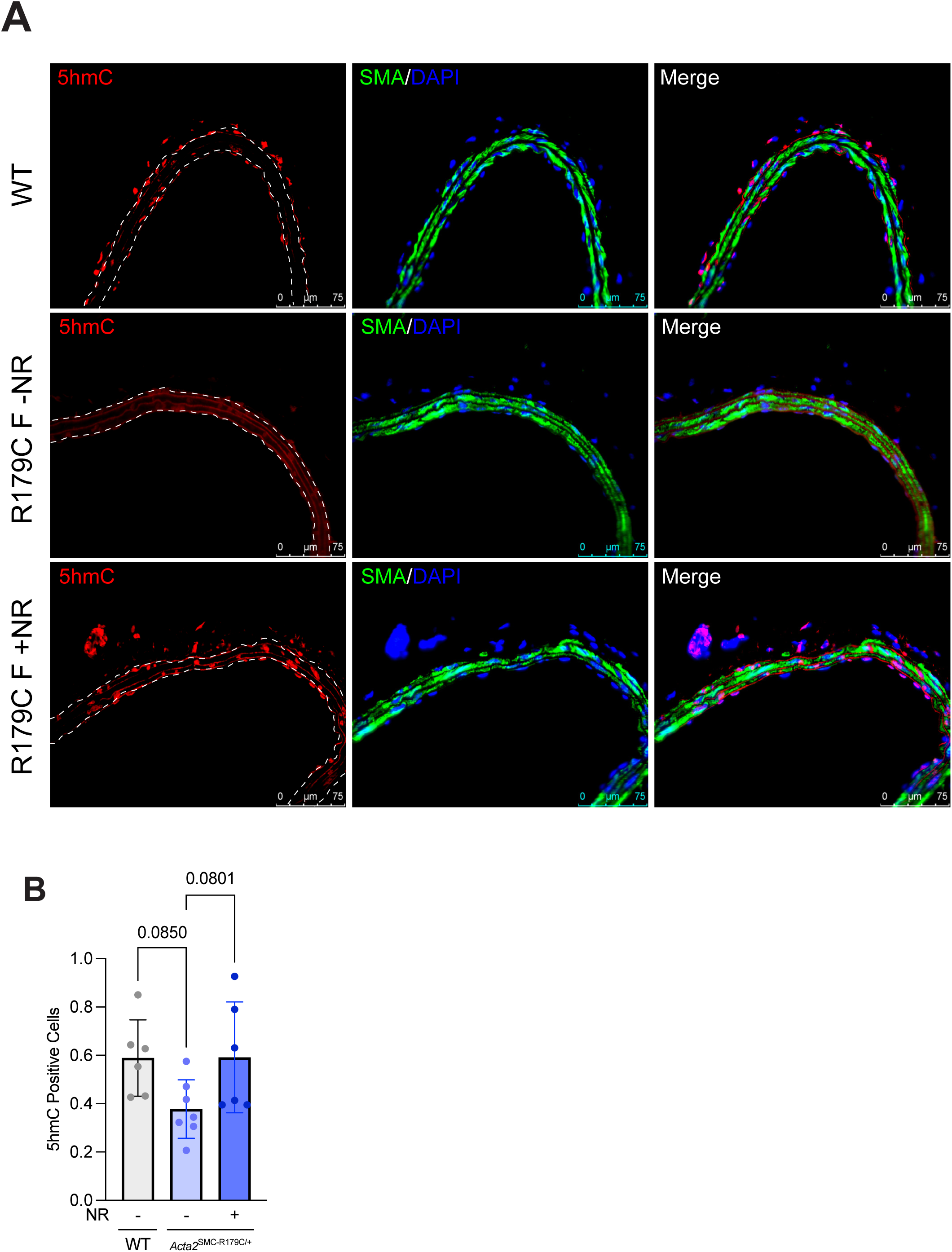
*Acta2*^SMC-R179C/+^ mice have decreased 5hmC in contralateral uninjured carotid artery tissue *in vivo* after left carotid artery ligation, increased by NR treatment. **A,B** Representative 5hmC immunostaining (red) in the contralateral uninjured right carotid artery at postoperative day 21 (A). Mutant tissue shows lower staining and NR-treated mutant tissue shows higher staining, but the quantified group differences (B) are trending rather than statistically significant. Green fluorescence shows α-SMA stain and blue fluorescence shows nuclei stained with DAPI. Scale bars, 75 µm. Each data point represents an individual mouse (n=7-8 per group). Statistical significance was assessed by two-way ANOVA with Tukey’s post hoc correction for multiple comparisons. P-values are indicated above brackets. All data are presented as mean ± standard deviation. SMC, smooth muscle cell; WT, wild-type; R179C, *Acta2*^R179C/+^; NR, nicotinamide riboside; α-SMA, α-smooth muscle actin; 5hmC, 5-hydroxymethylcytosine.

### Exogenous L-2 hydroxyglutarate (L-2-HG) prevents NR from inducing SMC differentiation

To test whether exogenous L-2-HG counteracts the effects of NR on contractile marker expression, we treated WT and *Acta2*^R179C/+^ SMCs with both NR and L-2-HG. If our mechanism is correct, exogenous 2-HG should prevent the rescue of SMC phenotype by NR. At both at a transcriptional and translational level, NR treatment alone induces an upregulation of contractile markers in both WT and *Acta2*^R179C/+^ SMCs. Conversely, exposure to L-2-HG results in reduced expression of these markers. After cotreatment for six days, contractile marker expression was lower than with NR alone in both genotypes, although the response varied by marker. These findings indicate that L-2-HG attenuates NR-induced contractile marker expression (Figure 4A,B, S2B).

**Fig. 4:**
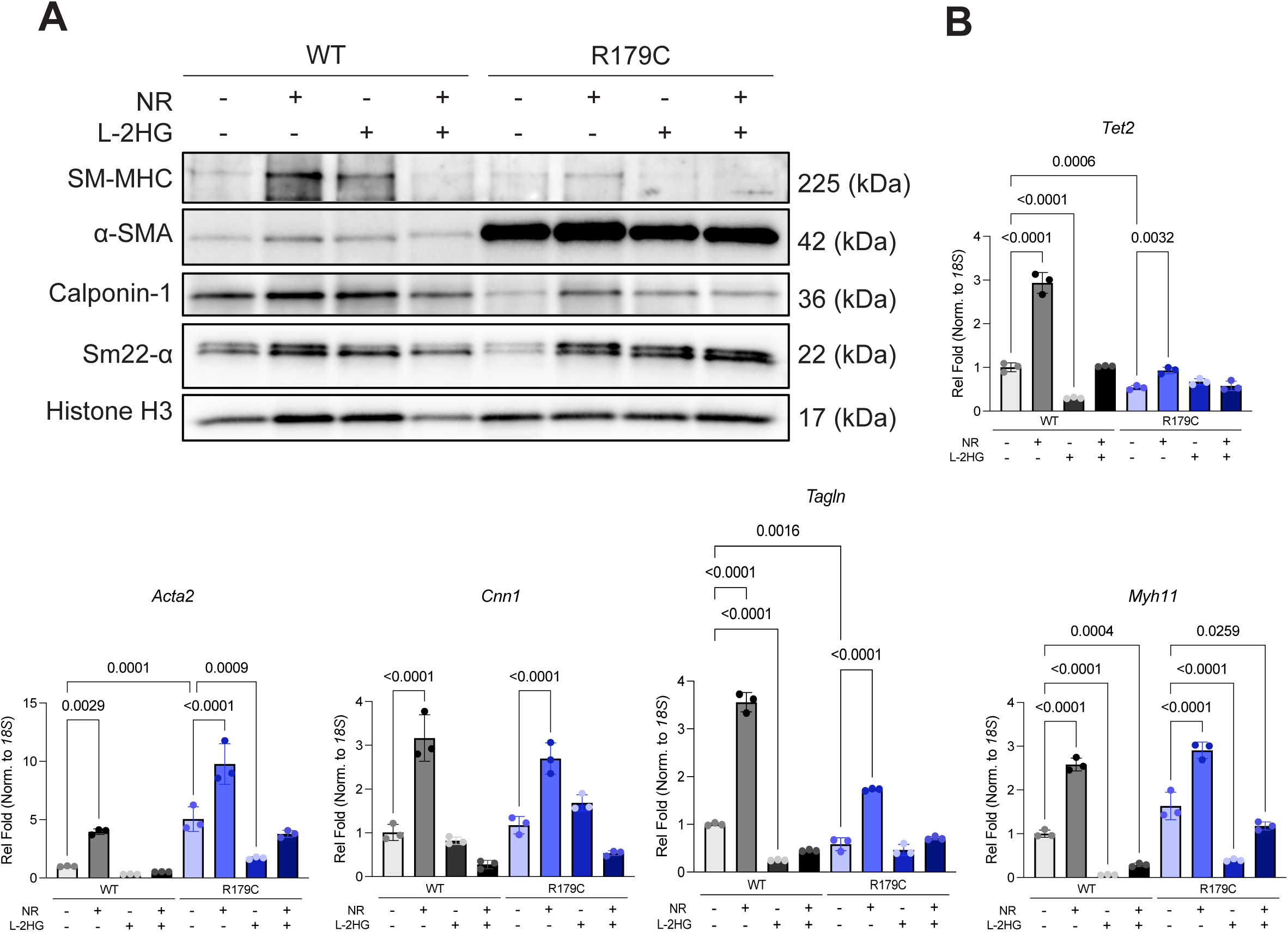
Exogenous L-2-HG prevents NR-mediated increase in contractile genes in *Acta2*^R179C/+^ SMCs. **A,B** Immunoblot analysis (A) demonstrates a trend toward increase levels of contractile proteins in *Acta2*^R179C/+^ SMCs after NR treatment, while co-treatment with L-2-HG (50 nM) prevents this trend. Gene expression shows the same trend assessed by RT-qPCR (B). Each data point represents a technical replicate (n=3), and results are representative of more than four independent experiments. Statistical significance was assessed by two-way ANOVA with Tukey’s post hoc correction for multiple comparisons. Quantifications of independent immunoblot replicates can be found in Supplemental Figure 2B. P-values are indicated above brackets. All data are presented as mean ± standard deviation. SMC, smooth muscle cell; TET2, ten-eleven translocation 2; WT, wild-type; R179C, *Acta2*^R179C/+^; NR, nicotinamide riboside; SM-MHC, smooth muscle myosin heavy chain; α-SMA, α-smooth muscle actin; L-2-HG, L-2-hydroxyglutarate.

### Exogenous L-2-HG de-differentiates WT SMCs via TET2 inhibition

To confirm that the 2-HG-TET2 axis is involved in regulation of SMC phenotype in WT SMCs, we treated WT SMCs with increasing concentrations of L-2-HG. Increasing L-2-HG concentrations reduced contractile marker transcripts. Representative immunoblots also showed lower contractile protein levels at higher doses (Figure 5A, B, S3A). L-2-HG reduced *Tet2* mRNA, but the independent immunoblot quantification did not show a significant reduction in TET2 protein (Figure 5A, B, S3A). D-2-HG also reduced contractile marker expression, (Supplemental Figure S3B,C). Addition of L-2-HG also induces a metabolic switch in WT SMCs: extracellular flux analyses demonstrated that exogenous L-2-HG significantly decreases the oxygen consumption rate (OCR) (Figure 5C, S4A) and significantly increases the extracellular acidification rate (ECAR) of WT SMCs (Figure 5D, S4B). This induced metabolic switch closely resembles the metabolic profile of *Acta2*^R179C/+^ SMCs, in which glycolysis is strongly favored over OXPHOS for energy production^17^. To assess the accompanying epigenetic changes, we measured 5hmC recovery, global 5mC levels, and TET2 chromatin occupancy after L-2-HG treatment. As expected, global 5hmC enrichment is significantly decreased with the addition of exogenous L-2-HG, and global 5mC enrichment is significantly increased (Figure 5F,G, S4C). Evidence of site-specific changes in TET2-DNA interaction were observed using ChIP in WT SMCs following the addition of L-2-HG. A significant decrease in TET2 abundance was observed within CArG boxes of key contractile SMC gene promoter regions (*Myh11*, *Acta2*, *Cnn1*, *Tagln*, and *Tspan2)* (Figure 5E).

**Fig. 5:**
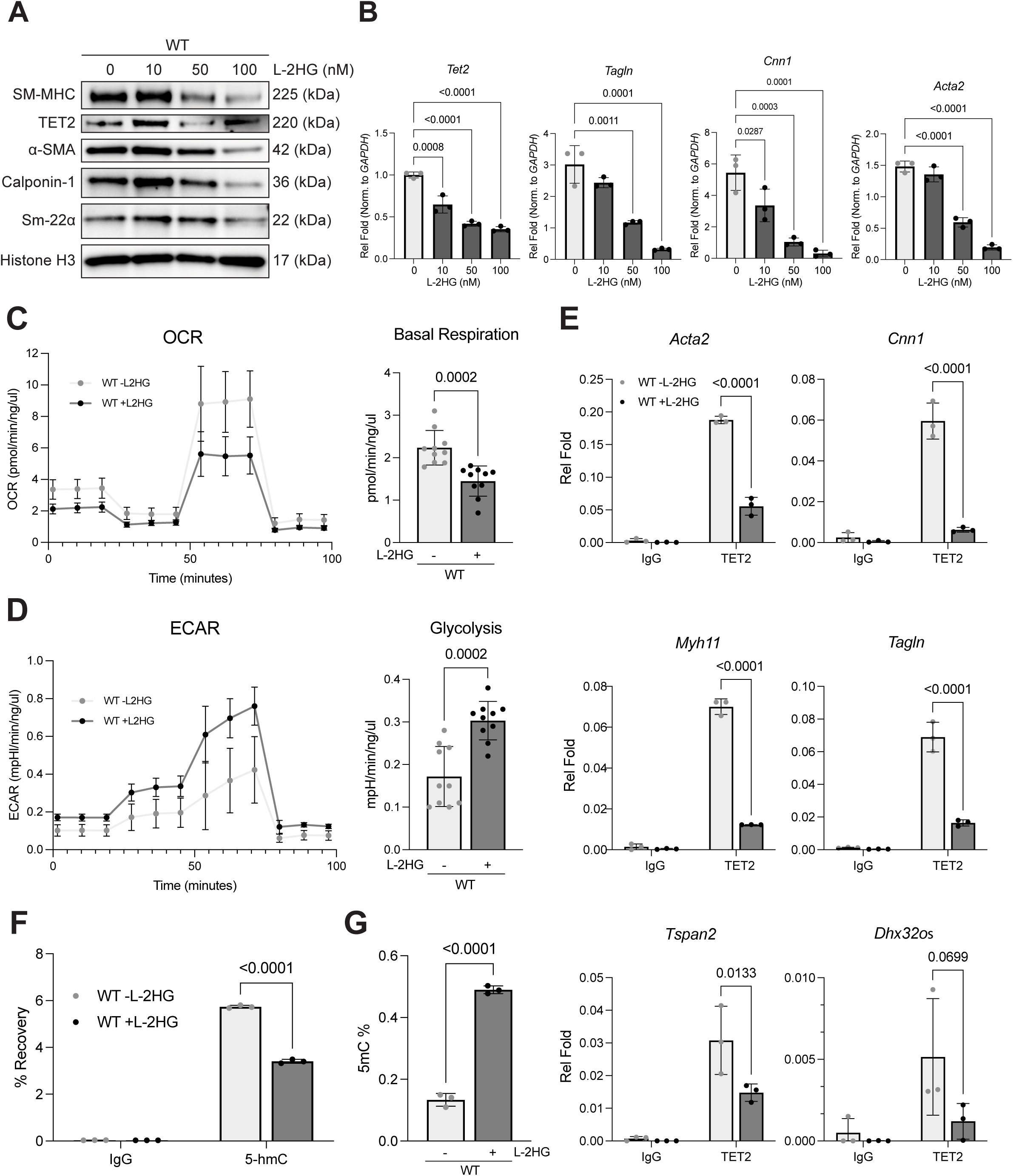
Exogenous L-2-HG reduces contractile marker expression, TET2 occupancy, and DNA hydroxymethylation in WT SMCs. **A,** Immunoblot analysis of contractile markers (SM-MHC, α-SMA, Calponin-1, Sm-22α) and TET2 in WT SMCs treated with increasing concentrations of L-2-HG (0, 10, 50, and 100 µM). Representative immunoblots show lower contractile protein levels at higher L-2-HG doses. Independent immunoblot quantification is shown in Supplemental Figure S3A, Histone H3 was used as a loading control. **B,** RT-qPCR analysis of *Tet2*, *Tagln*, *Cnn1*, and *Acta2* expression in WT SMCs showing reduced expression with increasing concentrations of L-2-HG (0, 10, 50, and 100 µM). Expression is normalized to *18S* and presented as relative fold change. Statistics were performed using one-way ANOVA with Dunnett’s multiple comparisons test, comparing each dose to the untreated (0 µM) control. **C, D,** Seahorse XF extracellular flux analysis of WT SMCs treated with or without 50 µM L-2-HG, measuring oxygen consumption rate (OCR) as a readout of mitochondrial respiration (C) and extracellular acidification rate (ECAR) as a readout of glycolysis (D). Quantification of basal respiration (C, right) and glycolysis (D, right) shows that L-2-HG shifts WT SMCs away from OXPHOS and toward glycolysis. Statistics were performed using an unpaired t-test. **E,** Chromatin immunoprecipitation (ChIP) analysis of TET2 occupancy at CArG-containing promoter regions of SMC contractile genes (*Tagln*, *Dhx32os*, *Cnn1*, *Acta2*, *Myh11*, and *Tspan2*) in WT SMCs treated with or without 50 µM L-2-HG shows reduced TET2 occupancy at five loci. Statistics were performed using an unpaired t-test. **F,G**, Global 5-hydroxymethylcytosine (5hmC) enrichment was assessed by hydroxymethylated DNA immunoprecipitation (hMeDIP), presented as percent recovery (F), and global 5-methylcytosine (5mC) levels quantified by MethylFlash ELISA, presented as 5mC percentage of total DNA (G), in WT SMCs treated with or without 50 µM L-2-HG. L-2-HG decreases global 5hmC and increases global 5mC. Statistics were performed using an unpaired t-test. For all bar graphs in this figure, each dot represents a technical replicate, and the data are representative of at least three independent experiments. All data are presented as mean ± standard deviation. SMC, smooth muscle cell; L-2-HG, L-2-hydroxyglutarate; TET2, ten-eleven translocation 2; OCR, oxygen consumption rate; ECAR, extracellular acidification rate; hMeDIP, hydroxymethylated DNA immunoprecipitation; ChIP, chromatin immunoprecipitation; 5hmC, 5-hydroxymethylcytosine; 5mC, 5-methylcytosine; WT, wild-type.

### Endogenous accumulation of L-2-HG dedifferentiates WT cells

L2HGDH is the only enzyme known to metabolize L-2-HG. This L-2-HG specific dehydrogenase can convert by oxidation L-2-HG into 2-Oxogluterate (2-OG), also known as α-ketoglutarate. WT SMCs treated with shRNAs targeting *L2HGDH* saw a significant reduction in the expression of *L2hgdh* by RT-qPCR. Only 2 out of the 5 shRNA(s) had a knockdown efficiency greater than 50% (Figure 6A, data not shown). Specific chiral separation and quantitation of L-2-HG by LC-MS/MS confirmed the endogenous accumulation of L-2-HG in WT SMCs after *L2HGDH* knockdown with construct #34 (Figure 6B). This endogenous accumulation of L-2-HG via *L2hgdh* knockdown resulted in the dedifferentiation of WT SMCs, with a reduction in the protein levels of key contractile markers like SM-MHC, α-SMA, and SM22α (Figure 6C, S4D). Together, these findings demonstrate that increased L-2-HG, whether by exogenous addition or increased endogenous accumulation, reduces the expression and activity of TET2, thereby inducing dedifferentiation of WT SMCs through the epigenetic silencing of contractile gene expression.

**Fig. 6:**
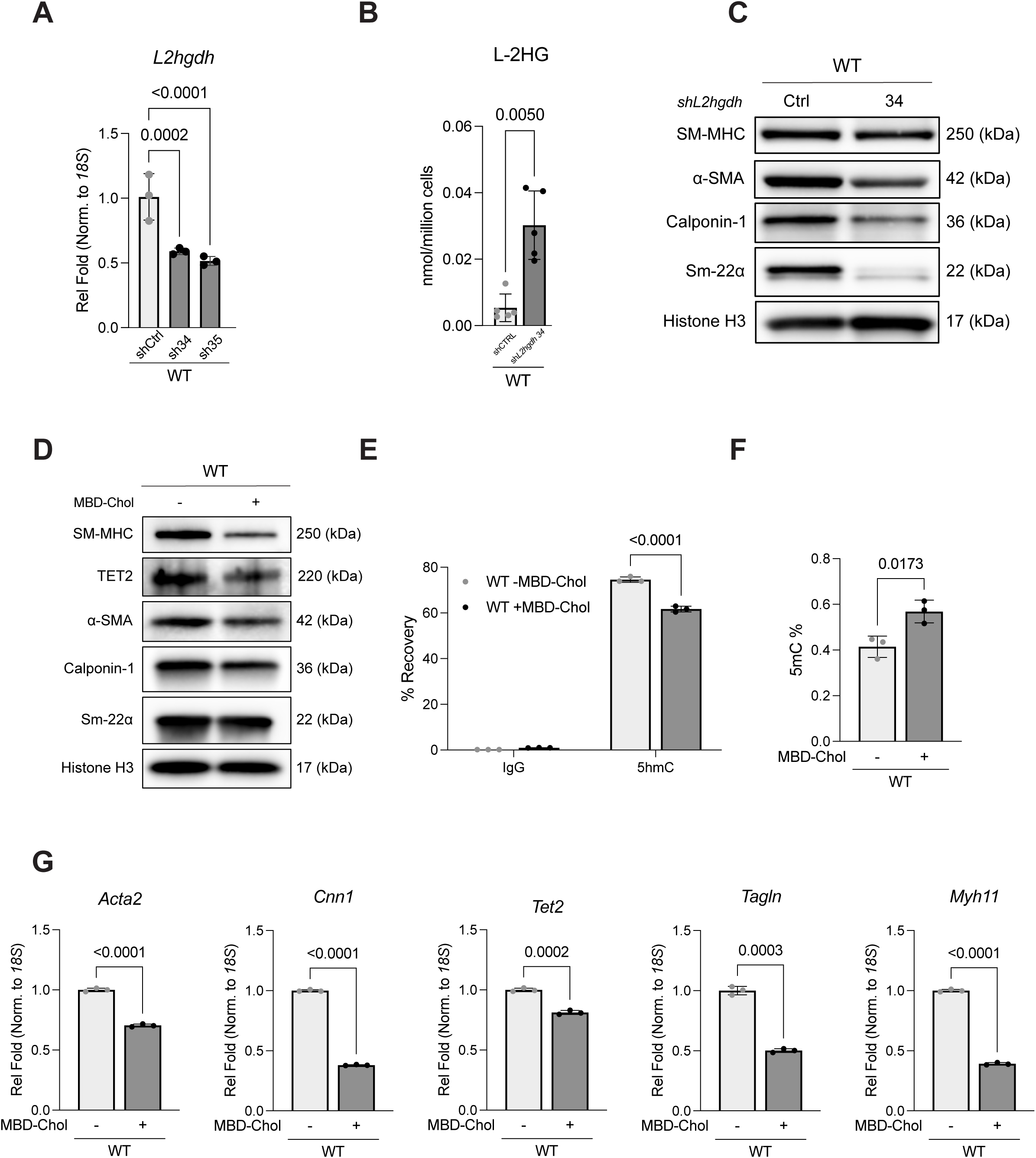
Endogenous accumulation of L-2HG and Cholesterol treatment induces dedifferentiation of WT SMCs. **A** Transient transfection of WT SMCs with two distinct shRNA constructs for *L2HGDH* showed a marked reduction in *L2hgdh* of ∼50%. **B,** LC/MS quantification of L-2-HG demonstrated that lentiviral knockdown of *L2hgdh* using construct 34 in WT SMCs resulted in a significant accumulation of L-2-HG. **C,** Immunoblot of WT SMCs control or sh*L2hgdh* construct 34 showed a reduction of contractile markers after *L2hgdh* knockdown. **D,G,** Treatment with 10ng/mL MBD-Chol for 72 hours decreases accumulation of SMC contractile proteins and TET2 by immunoblot (D) and concomitantly decreases expression of the genes by RT-qPCR (G). **E,F,** Global 5-hydroxymethylcytosine (5hmC) enrichment was assessed by hydroxymethylated DNA immunoprecipitation (hMeDIP), presented as percent recovery (E), and global 5-methylcytosine (5mC) levels quantified by MethylFlash ELISA, presented as 5mC percentage of total DNA (F), in WT SMCs treated with or without MBD-Chol. MBD-Chol decreases global 5hmC and increases global 5mC. Statistical significance was assessed for two groups (B-G) using an unpaired two-tailed t-test with Welch’s correction and for three or more groups (A) by One-way ANOVA with Tukey’s post hoc correction for multiple comparisons. Quantifications of independent immunoblot replicates can be found in Supplemental Figure 4D,E. P-values are indicated above brackets. All data are presented as mean ± standard deviation. SMC, smooth muscle cell; TET2, ten-eleven translocation 2; OCR, oxygen consumption rate; ECAR, extracellular acidification rate; hMeDIP, hydroxymethylated DNA immunoprecipitation;; 5hmC, 5-hydroxymethylcytosine; 5mC, 5-methylcytosine; WT, wild-type.

### Cholesterol-induced phenotypic modulation is accompanied by reduced *TET2* expression and DNA hydroxymethylation

Treatment with MBD-Chol has been previously shown to reduce expression of SMC contractile genes, along with upregulation of markers of atherosclerosis-associated phenotypic modulation^24^. SMC phenotypic modulation is also accompanied by a metabolic shift from OXPHOS to glycolysis^8,31^. However, the mechanisms linking cholesterol-induced metabolic and transcriptional changes are unknown. To determine whether cholesterol-induced phenotypic modulation is accompanied by changes in TET2 and DNA hydroxymethylation, we treated cultured WT SMCs with 10 µg/mL MBD-Chol for 72 hours. In accordance with published results, contractile gene expression and protein levels are significantly decreased, as are levels of TET2 (Figure 6D, G, S4E). Global levels of 5hmC are decreased (Figure 6E) and global levels of 5mC are increased following MBD-Chol treatment(Figure 6F, S4F), supporting that TET2 activity is globally inhibited. These findings associate cholesterol-induced phenotypic modulation with reduced TET2 expression and altered DNA methylation and hydroxymethylation.

## Discussion

Our data support the 2-HG-TET2 axis as a mechanistic link between metabolic state and SMC differentiation. In *Acta2*^R179C/+^SMCs, elevated 2-HG levels coincide with reduced TET2 expression, diminished 5hmC levels, and loss of TET2 occupancy at CArG-containing contractile gene promoters. These defects are rescued by the NR-mediated metabolic shift towards OXPHOS, although TET2 occupancy was not significantly restored at every locus. Critically, both exogenous addition and endogenous accumulation of 2-HG are sufficient to dedifferentiate WT SMCs, recapitulating the metabolic and epigenetic features of *Acta2*^R179C/+^SMCs. These results confirm that 2-HG acts as a metabolic determinant of SMC dedifferentiation independent of the *ACTA2* variant. Further, cholesterol, a stimulus known to drive SMC phenotypic modulation^24^, reduces TET2 expression and 5hmC recovery in WT SMCs. Whether cholesterol causes these changes through increased 2-HG remains to be determined.

TET2 has been well established as a key regulator of SMC differentiation, in part through its cooperation with chromatin-modifying enzymes—such as the histone acetyltransferase p300—to promote a contractile gene program^22,23^. TET2 localizes to CArG-rich regulatory regions within key SMC transcriptional programs, including the promoters of *MYOCD* and *SRF*, where it facilitates transcriptional activation of contractile genes^22,32^. Under physiological conditions, myocardin (MYOCD) associates with serum response factor (SRF) to form a master regulatory complex that drives SMC differentiation while simultaneously repressing transcription factors such as *KLF4* that promote dedifferentiation^22,33–35^. Cholesterol treatment induces KLF4 expression while simultaneously downregulating TET2; TET2’s established role as a potent inhibitor of KLF4 likely explains this observation^24,36^. Our ChIP data indicate that TET2 occupancy at several CArG-containing SMC loci is reduced in *Acta2^R^*^179^*^C/+^* SMCs and increases at a subset of loci with NR treatment. These observations are consistent with metabolic regulation of TET2-associated chromatin changes. Given that dedifferentiated SMCs, such as those found in atherosclerotic plaques, exhibit a similar metabolic profile of increased glycolysis and decreased OXPHOS^8,37–39,^ 2-HG-driven suppression of TET2 may represent a candidate mechanism linking metabolic rewiring with epigenetic silencing of the contractile program across vascular disease contexts.

While our data strongly support the 2-HG-TET2 axis as a potential mediator of SMC phenotypic modulation, 2-HG is not exclusively selective for TET2; rather it broadly targets α-KG–dependent dioxygenases, suggesting that additional epigenetic mechanisms may contribute to the observed phenotypes^19^. Although TET2 is the most extensively characterized TET enzyme in SMC biology, both TET1 and TET3 have also been implicated in SMC plasticity. TET1 has been associated with promoting the transition from a contractile to a synthetic, proliferative phenotype during atherosclerosis^40^, and TET3 has been linked to pathogenic SMC responses under inflammatory conditions^41^. Given that all TET enzymes are α-KG–dependent dioxygenases, it remains unclear to what extent 2-HG accumulation impacts TET1 and TET3 activity within our model system. Additionally, 2-HG is a known inhibitor of Jumonji C (JmjC) domain-containing histone demethylases (KDMs), which regulate chromatin accessibility by altering histone methylation, including H3K9 and H3K36 methylation^19,42–44^. Inhibition of these enzymes by 2-HG could reinforce a transcriptionally repressive chromatin state at SMC differentiation loci, thereby acting in parallel with TET2 suppression. While our findings establish that 2-HG accumulation in SMCs is sufficient to suppress TET2 activity and drive phenotypic modulation, we cannot discount broader epigenetic effects through coordinated inhibition of multiple α-KG–dependent enzymes. TET2 loss-of-function studies, combined with rescue using wild-type or catalytically inactive TET2, would help distinguish TET2-dependent effects from those involving other α-KG-dependent enzymes. Therefore, a comprehensive evaluation of histone modifications and dioxygenase targets will be necessary to fully delineate the epigenetic effects of 2-HG on the SMC epigenome.

There are two enantiomers of 2-HG, both of which inhibit α-ketoglutarate-dependent dioxygenases, including TET2^19,42^. Oncogenic mutations in *IDH1* and *IDH2* are characterized by the aberrant accumulation of D-2-hydroxyglutarate (D-2-HG), which functions as a prototypical oncometabolite and is prevalent in malignancies such as gliomas, acute myeloid leukemia, and cholangiocarcinoma^45–47^. By contrast, the L-2-HG enantiomer is produced by noncanonical activities of lactate dehydrogenase and malate dehydrogenase in the context of hypoxic or acidic environments^48,49^. Elevated levels of either D- or L-2-HG promote widespread DNA and histone hypermethylation through inhibition of α-KG–dependent dioxygenases, establishing a differentiation blockade that contributes to tumor initiation and progression^19,42,50^. Importantly, we confirmed that both D- and L-2-HG drive dedifferentiation of WT SMCs via inhibition of TET2 (Figure 5A,B S3A,B,C).

Our findings suggest that metabolic reprogramming—specifically promoting OXPHOS over glycolysis—can effectively reduce 2-HG accumulation and restore epigenetic balance. In SMCs, this shift is sufficient to rescue TET2 activity and reestablish a contractile phenotype, highlighting the potential of targeting metabolic flux as a therapeutic strategy to promote SMC identity and vascular integrity. While our data focused on NR, alternative metabolites such as L-ascorbic acid have been shown to enhance TET enzyme activity and promote DNA demethylation, in part through redox regulation and support of mitochondrial metabolism, thus restoring differentiation programs in certain cellular contexts^51^. More broadly, these observations underscore the therapeutic potential of targeting the interface between metabolism and epigenetic regulation to correct aberrant cell fate decisions. Such strategies may prove particularly valuable in diseases characterized by metabolic rewiring and epigenetic instability, where restoring physiological metabolite levels could reverse pathogenic gene expression programs.

### What Are the Clinical Implications?

Vascular smooth muscle cell (SMC) phenotypic modulation, i.e., the loss of contractile identity, drives atherosclerosis, aortic aneurysm, and moyamoya disease, yet no therapies target the underlying metabolic reprogramming. This study identifies the oncometabolite 2-hydroxyglutarate (2-HG) as a link between the glycolytic state of dedifferentiated SMCs and epigenetic silencing of the contractile program through inhibition of TET2 and loss of DNA hydroxymethylation. Both exogenous and endogenous 2-HG accumulation are sufficient to dedifferentiate wild-type SMCs, and boosting oxidative metabolism with nicotinamide riboside (NR) lowers 2-HG and restores markers of SMC identity. These findings provide a rationale for metabolic interventions aimed at preserving SMC differentiation in vascular disease. Because this work was performed in mouse SMCs and does not establish that TET2 is required for the effects of NR, validation in human SMCs, direct tests of TET2 dependence, and in vivo disease-outcome studies are needed before clinical translation.

## Author Contributions

Conceptualization: C.S.K., Methodology: J.E.E.P., A.Ro., A.D., A.C., E.G., Investigation and data analysis: J.E.E.P., A.Ro., A.Ri., A.D., J.G.S., A.D.G., H.K., J.C., M.C., K.E.B., S.S.R., A.C., C.S.K.; Supervision: D.M.M., C.S.K., Funding Acquisition: A.C., D.M.M., C.S.K., Writing – original draft preparation: J.E.E.P., C.S.K.; Writing – review & editing: all authors

## Non-standard Abbreviations and Acronyms

2-HG: 2-hydroxyglutarate
5hmC: 5-hydroxymethylcytosine
5mC: 5-methylcytosine
α-KG: α-ketoglutarate
ChIP: chromatin immunoprecipitation
ECAR: extracellular acidification rate
hMeDIP: hydroxymethylated DNA immunoprecipitation
LCAL: left carotid artery ligation
MBD-Chol: methyl-β-cyclodextrin-complexed cholesterol
MMD: moyamoya disease
MSMDS: multisystemic smooth muscle dysfunction syndrome
NR: nicotinamide riboside
OCR: oxygen consumption rate
OXPHOS: oxidative phosphorylation
POD: postoperative day
SMC: smooth muscle cell
TET2: ten-eleven translocation 2
WT: wild-type

---

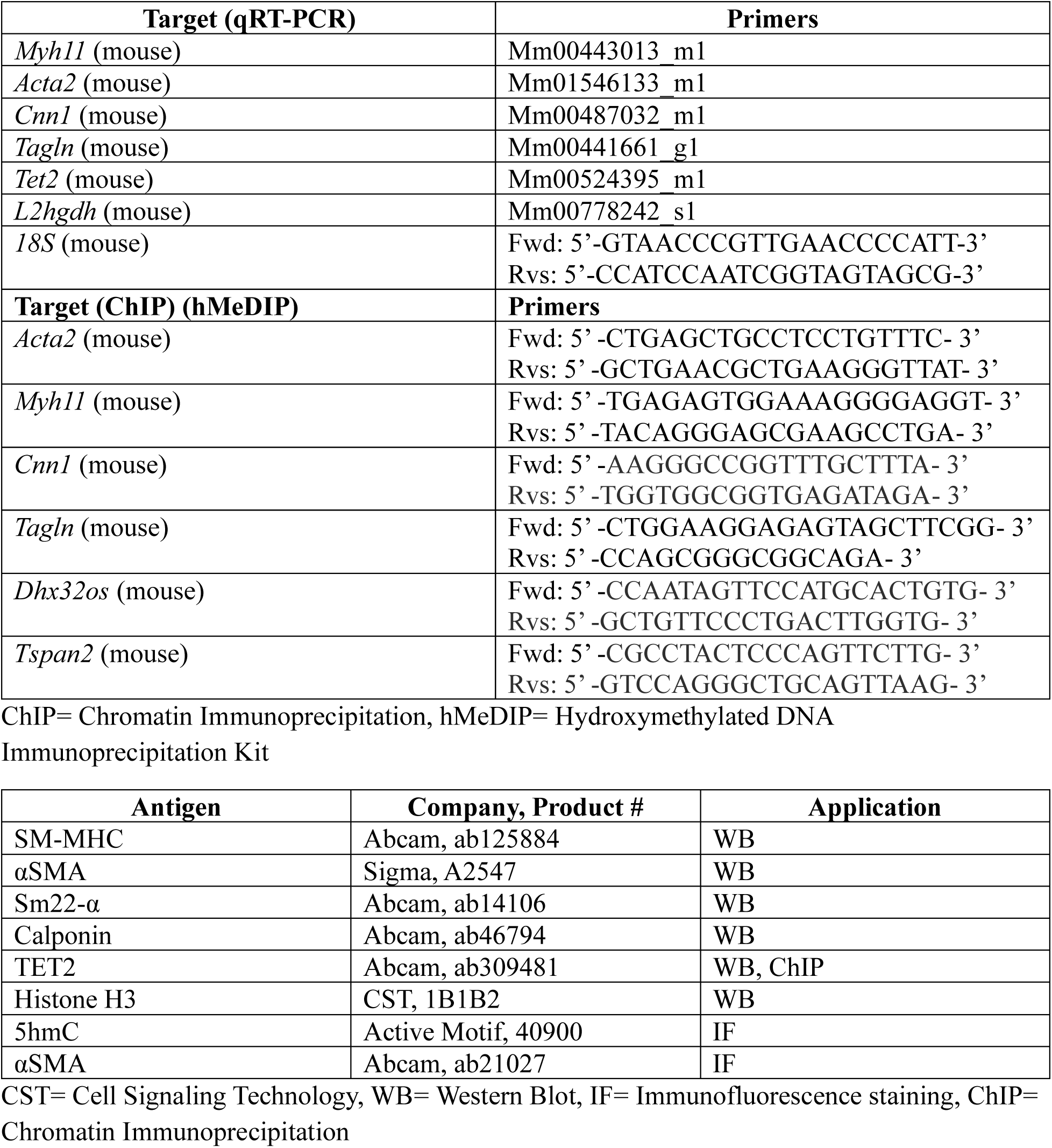

