## Supplemental Figures for "2-hydroxyglutarate-dependent inhibition of TET2 is a molecular link between metabolic reprogramming and smooth muscle cell differentiation"

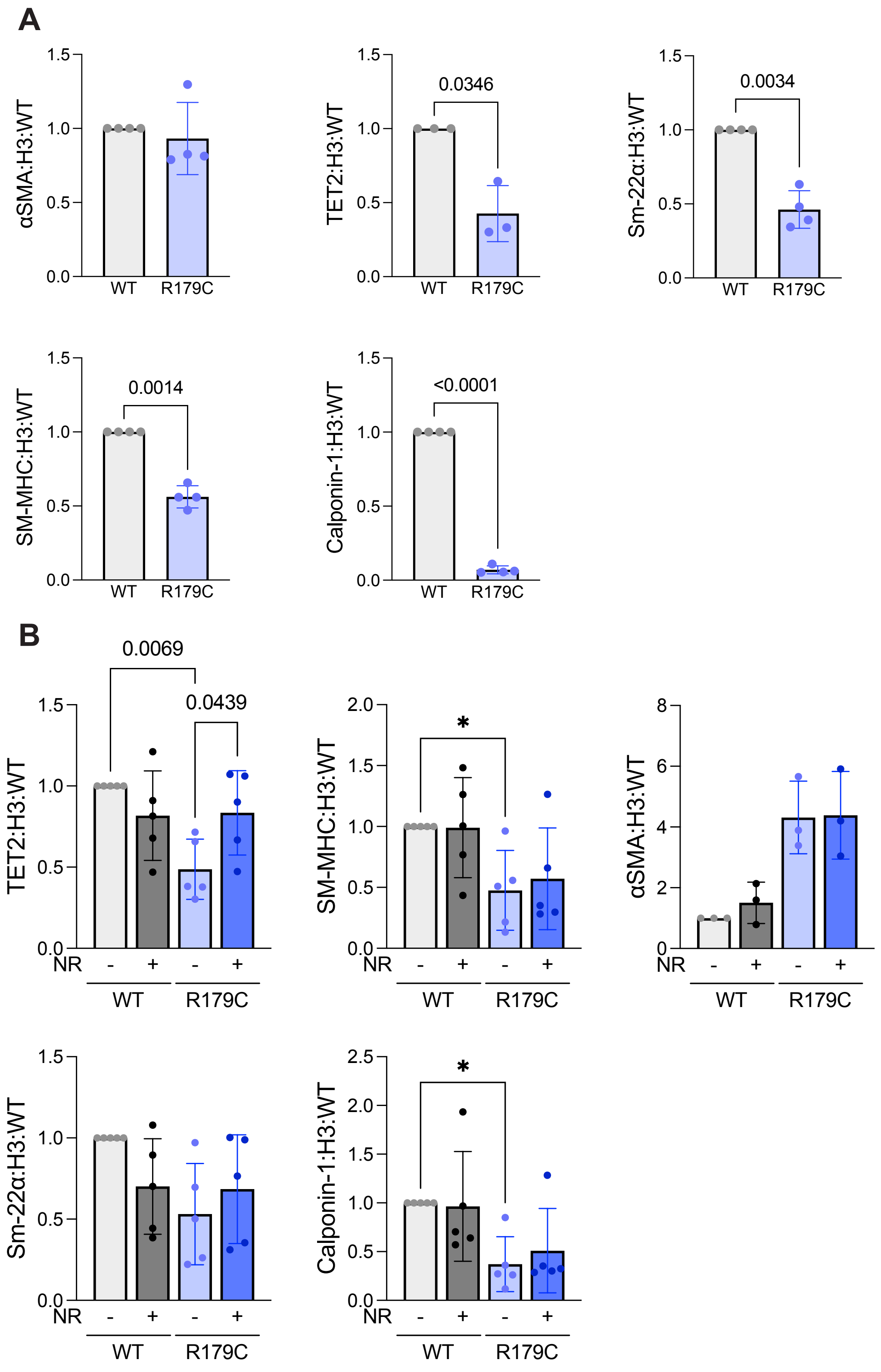


**Figure S1. A.** Western blot quantification for Figure 1A. Each dot represents an independent experimental replicate (n=4). **B.** Western Blot quantification for Figure 1C. Each dot represetns an independent experimental replicate (n=5). For all graphs in this figure, statistics were performed via unpaired students' t-test (for two sample comparisons S1A) and Repeated Measures two-way ANOVA with Šídák's multiple comparison test (S1B).

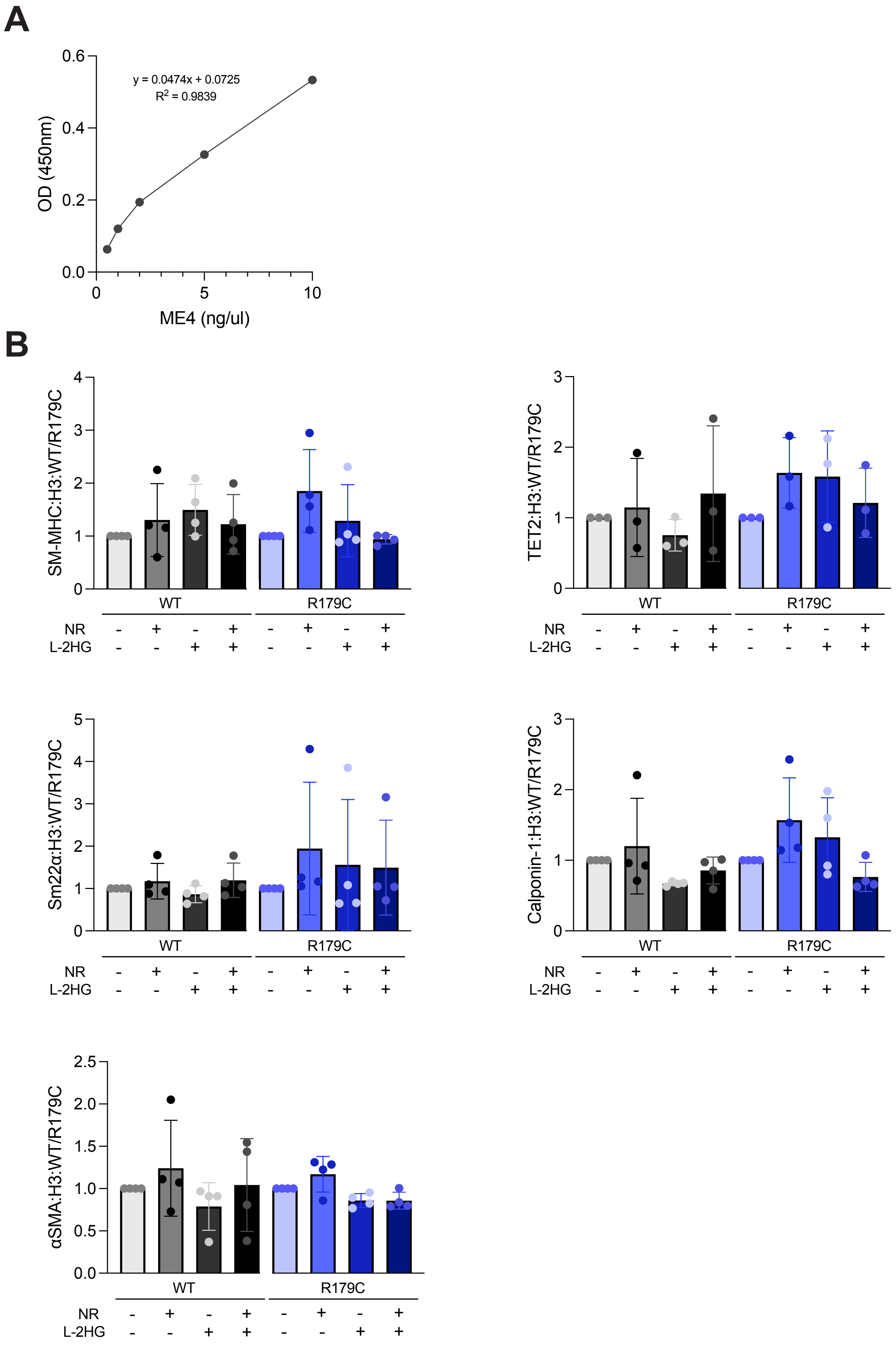


**Figure S2. A.** Standard curve generated for analysis of the 5mC ELISA assay shown in Figure 2B. **B.** Western blot quantification for Figure 3A. Each dot represents an independent experiemental replicate (n=4). For all graphs in this figure, statistics were performed via Reapeated Measures two-way ANOVA with Šídák's multiple comparison test (S2B).

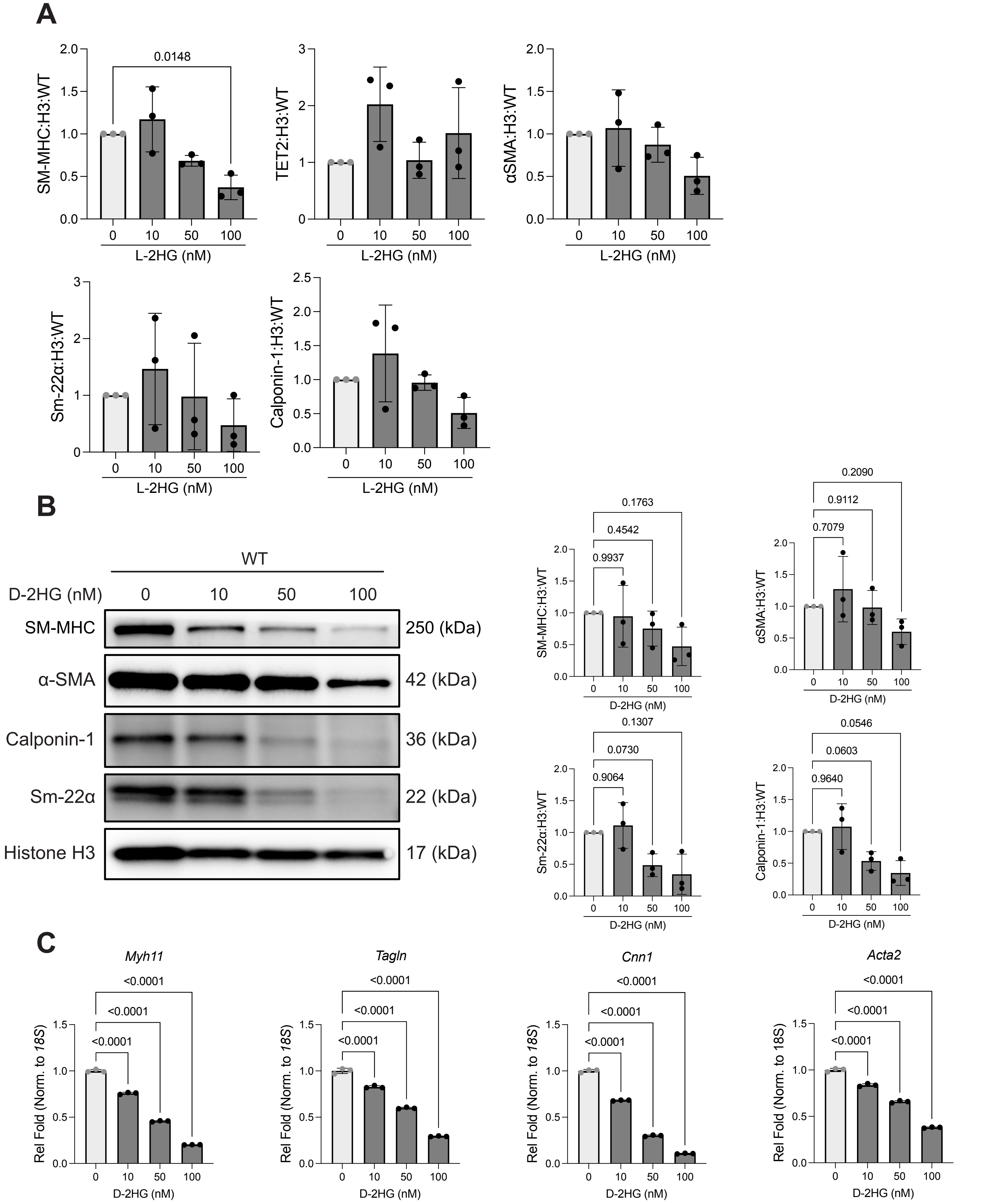


**Figure S3. A.** Western blot quantification for Figure 5A (n=3). **B.** Western Blot showing dose-dependent dedifferentiation of WT SMCs after 72 hr treatment of D-2-Hydroxygluterate (D-2HG). Immunoblot is a representative of at least 3 independent replicates (n=3). Western Blot Quantification of S3B (shown right). Each dot represents an independent experimental replicate (n=3). **C.** RT-qPCR of WT SMCs after 72 hr treatment of D-2HG for contractile genes (*Acta2, Tagln, Myh11, Tet2, Cnn1*). For all graphs in this figure, statistics were performed via Repeated Measures one-way ANOVA with Šídák's multiple comparison test (S3A, S3B), and one-way ANOVA with Turkey's multiple comparisons test (S3B).

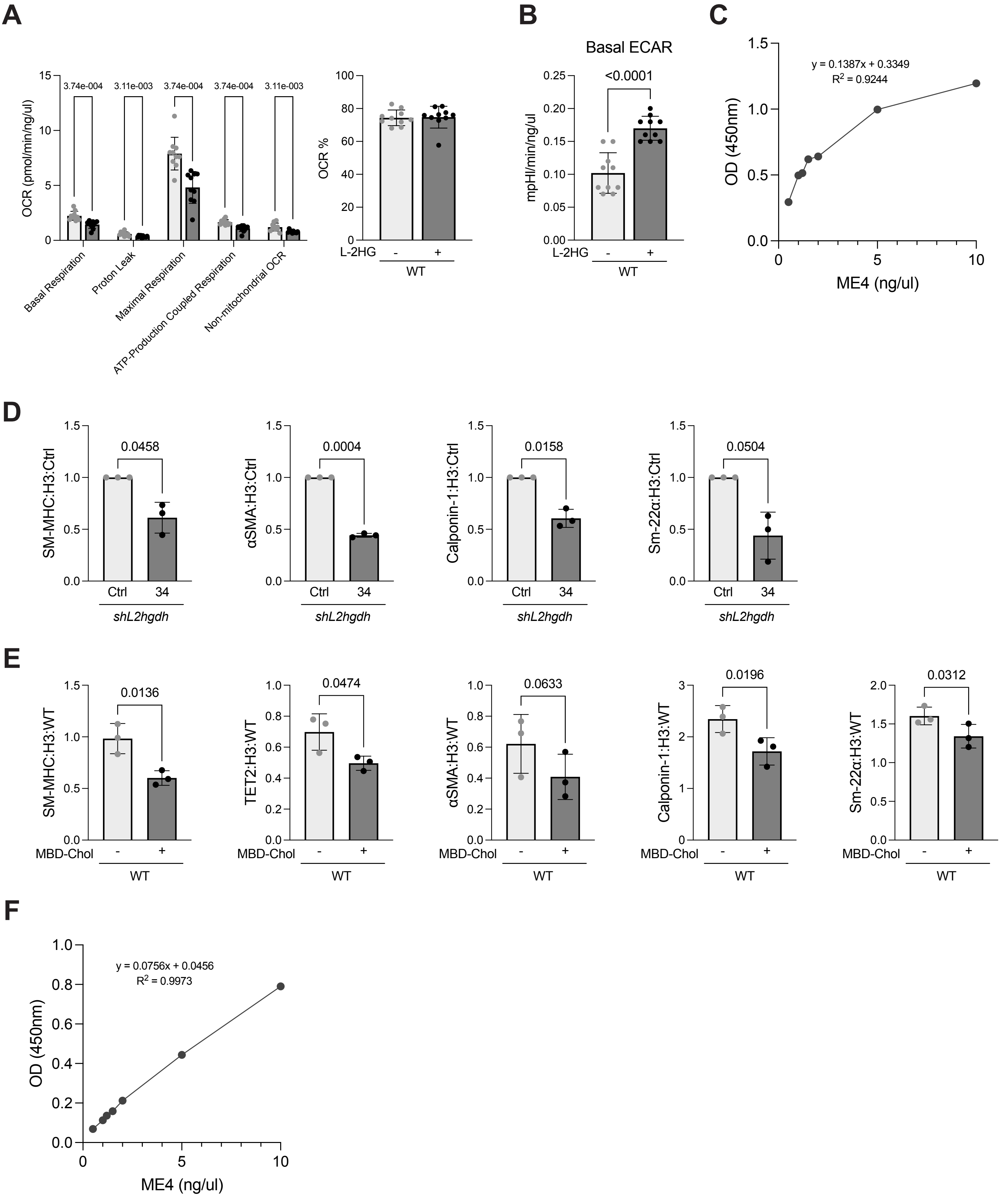


**Figure S4 A.** Individual parameters from the Mito Stress Test Seahorse Analysis shown in Figure 5C. **B.** Basal extracellular acidification rate from the Glycolysis Stress Test Seahorse Analysis shown in Figure 5D. For S4A-B, each dot represents a technical replicate from a representative experiment of at least 3 independent experiments (n=10). **C & F.** Standard curve generated for analysis of the 5hmC ELISA assay shown in Figure 5G and Figure 6D. **D.** Western Blot quantification from Figure 6B, each dot represents an independent experimental replicate (n=3). **E.** Western Blot quantification from Figure 6D, each dot represents an independent experimental replicate (n=3). For all graphs in this figure, statistics were performed via students' t-test (For two sample comparisons SA-B, E-F)
